# An intact keratin network is crucial for mechanical integrity and barrier function in keratinocyte cell sheets

**DOI:** 10.1101/661462

**Authors:** Susanne Karsch, Fanny Büchau, Thomas M. Magin, Andreas Janshoff

**Affiliations:** Institute of Physical Chemistry, University of Göttingen, Germany; Institute of Biology and SIKT, University of Leipzig, Germany

## Abstract

The isotype-specific composition of the keratin cytoskeleton is important for strong adhesion, force resilience, and barrier function of the epidermis. However, the mechanisms by which keratins regulate these functions are still incompletely understood. In this study, the role and significance of the keratin network for mechanical integrity, force transmission, and barrier formation were analyzed in murine keratinocytes. Following the time-course of single-cell wounding, wildtype (WT) cells slowly closed the gap in a collective fashion involving tightly connected neighboring cells. In contrast, the mechanical response of neighboring cells was compromised in keratin-deficient cells, causing an increased wound area initially and an inefficient overall wound closure. Furthermore, the loss of the keratin network led to impaired, fragmented cell-cell junctions and triggered a profound change in the overall cellular actomyosin architecture. Electrical cell-substrate impedance sensing of cell junctions revealed a dysfunctional barrier in knockout (Kty^−/−^) compared to WT cells. These findings demonstrate that Kty^−/−^ cells display a novel phenotype characterized by loss of mechanocoupling and failure to form a functional barrier. Re-expression of K5/K14 rescued the barrier defect to a significant extent and reestablished the mechanocoupling with remaining discrepancies likely due to the low abundance of keratins in that setting. Our study reveals the major role of the keratin network for mechanical homeostasis and barrier functionality in keratinocyte layers.

## Introduction

The mammalian epidermis is a multilayered barrier-forming epithelium that protects the body against physical and chemical insults and against dehydration. Its physico-chemical properties depend on the presence of adherens junctions, desmosomes, and tight junctions, which couple to the actin and keratin cytoskeletons through cadherin and zonula occludens family members, respectively^1^. Studies in model organisms and cultured mammalian cells have established an instructive role of E-cadherin in the formation of adherens junctions, desmosomes, and tight junctions, which depends on its ability to sense mechanical forces and transduce tension across cells in an actomyosin-dependent manner^2^. In the epidermis, where E-cadherin is present in all viable layers, its loss interferes with the tight junction barrier and resulted in perinatal death^3^.

In addition to the actin cytoskeleton associated with adherens and tight junctions, keratins (K) represent major cytoskeletal components of keratinocytes. In basal keratinocytes, K5 and K14 assemble through heterodimers into a three-dimensional keratin network anchored at desmosomes and hemidesmosomes to render the epidermis resilient against mechanical and other forms of stress^4^. During wound healing, a condition that requires a transient decrease in intercellular adhesion to promote enhanced migration and proliferation of keratinocytes for wound closure, keratins K6, K16, and K17 are transiently upregulated to form a more dynamic keratin network^5–7^. Both change of expression or deletion of all keratins render keratinocytes softer and more invasive in corresponding assays^8–10^. Furthermore, lack of keratins compromises desmosomal adhesion due to a decrease in desmoplakin, desmoglein 1, and plakophilin 1, accompanied by smaller and fewer desmosomes^10,11^. Impaired desmosome adhesion, observed in desmoplakin of Dsg-1 deficient cells and mice, not only affects mechanical integrity, but frequently resulted in alterations of actin-dependent processes, such as cell flattening and migration ^12–15^

Furthermore, there is emerging evidence for an involvement of desmosomes in the formation and function of epidermal tight junctions. Plakophilin 1-deficient mice suffered from disturbed tight junctions with an impaired inside-out barrier although expression of claudin-1, occludin, and ZO-1 was unaltered^16^, whereas expression of claudin-1 and transepithelial resistance were increased in desmoplakin-deficient cells^13^. In addition to an indirect effect of keratins on tight junctions via regulating desmosome composition, K76 was described to directly interact with claudin 1. The knockout of K76 was accompanied by mislocalization of the transmembrane protein claudin 1 and by a compromised tight junction barrier^17^. These observations raise the question whether keratins coordinate the keratinocyte response in an epithelial monolayer during wound closure, by coordinating the reorganization of the actin cytoskeleton, adherens and tight junctions^18,19^. Using single-cell wounding, we have previously shown that in MDCK II cells, healing of these micro-lesions was accompanied by an increase in pretension in cells neighboring the wound^20^.

Here, we have investigated the role of keratins in the process of gap closure by comparison of WT, keratin-deficient, and rescue keratinocytes re-expressing a single keratin pair. We find that upon single cell wounding, the mechanical response of keratin-deficient neighbors was completely abolished. Furthermore, lack of keratins caused a reorganization of the actin cytoskeleton, accompanied by mislocalization of adherens and tight junctions, leading to a TJ-associated barrier defect. Re-expression of keratins normalized the cortical actin cytoskeleton, adherens, and tight junctions, and partially restored the barrier. Our findings reveal a crucial role of keratins in mechanocoupling of keratinocytes during wound closure and in the correct localization and function of tight junction proteins.

## Materials & Methods

### Cell Culture

Generation of WT, keratin type I knock-out (Kty^−/−^) and rescue cells was carried out as described earlier^8,11,21^. Cultivation of keratinocytes was performed according to Seltmann et al.^22^ in FAD medium. Cells were seeded in low calcium medium for 24h; one day after seeding (48 h prior to the experiment), medium was changed to high calcium medium containing 1.2 mM CaCl_2_. For experiments without CO_2_ supply, 10 mM HEPES (Biochrom, Berlin, Germany) was added to the culture medium.

### Immunofluorescence analysis

In wounding experiments, cells were washed with PBS before fixation for 15 min in 4% formalin freshly prepared from paraformaldehyde (PFA, Fluka, Buchs, Switzerland) in PBS, and finally washed with PBS. Permeabilization of the membrane and blocking of unspecific binding was achieved by adding 5 % (w/v) bovine serum albumin (IgG-free, Carl Roth, Karlsruhe, Germany) and 0.3 % (w/v) Triton-X 100 (Sigma-Aldrich, St. Louis, USA) in PBS for 30 min at room temperature. Antibodies were diluted in 1 % (w/v) bovine serum albumin with 0.3 % (w/v) of Triton-X 100 in PBS and incubated for 1 h. Actin was stained with 165 nM Alexa Fluor 546 conjugated Phalloidin (Life Technologies, Carlsbad, USA) and keratins with 1 μg/ml anti-cytokeratin 14 (mouse monoclonal, Abcam, Cambridge, UK) and Alexa Fluor 488 IgG goat-anti mouse (Thermo Fisher Scientific, Waltham, USA) as secondary antibody. Confocal images were taken with the FluoView1200 system (Olympus, Tokyo, Japan) using an oil immersion objective (UPLFLN100xO2PH, NA 1.3; Olympus. Tokyo, Japan). Image analysis and processing were performed using the FluoView software (Olympus, Tokyo, Japan). For adherens or tight junction staining, cells were fixed for 30 min in ice-cold ethanol and 3 min in acetone at room temperature, followed by 3 washing steps with TBS. For actin and myosin staining, cells were fixed for 15 min in 4% formalin/PBS, made freshly from PFA, at 4°C, washed in TBS and then permeabilized for 5 min in 0.25% Triton/PBS. Antibodies were diluted in 1 % (w/v) bovine serum albumin in TBS and incubated for 1 h. Antibodies are listed in Supplementary table 1. Confocal images were taken with a Zeiss AxioImager equipped with Apotome2 using a 63 /1.4 NA oil immersion objectives. Image analysis and processing were performed using Zen Blue Software (Carl Zeiss, Inc.) and Photoshop CS4 (Adobe) software. LUT (lookup table; brightness and gamma) was adjusted using Photoshop.

### Western Blotting

SDS-PAGE was performed as described^21^. In short: total proteins were extracted in SDS-PAGE sample buffer under repeated heating (95°C) cycles. Separation of total protein extracts was carried out by standard procedures (8–10 % SDS-PAGE). Western blotting was performed as described earlier^21^. Antibodies are listed in Supplementary table 1. Statistical significance was determined by two-tailed t-tests (* indicating p <0.5, ** indicating p<0.01 and *** indicating p<0.001).

### Wound induction and optical analysis

Wounding was performed via a microinjection system (Femtojet and InjectMan NI2, both from Eppendorf, Hamburg, Germany) by moving a glass capillary (Femtotip; Eppendorf, Hamburg, Germany) across the cell body. The wounding was controlled with an inverted microscope (Olympus IX 81; Olympus, Tokyo, Japan) equipped with a 40x objective (LUCPLFKN 40XPH, NA = 0.6; Olympus, Tokyo, Japan). During wounding and video recording, the cells were placed in a petri dish heater set to 32 °C (JPK Instruments, Berlin, Germany). Videos during wounding were acquired with the cellSens Dimension software (Olympus, Tokyo, Japan) in live mode, resulting in 80 ms per frame. Phase contrast videos during wound closure were acquired with 1 frame/min. Cell size of the wounded cell was measured every 10 min during closure by manually selecting the cell boundaries. Image processing and analysis was done via ImageJ.

### Atomic force microscopy

Mechanical analysis after wounding was performed with force indentation experiments at the wounded spot on a NanoWizard II/IV atomic force microscope (AFM) (JPK Instruments, Berlin Germany). Cantilever calibration was done via the thermal noise method^23^ for every used cantilever (MLCT-C-cantilever from Bruker AFM Probes, Camarillo, USA). For individual force curves, performed on a grid with 2 μm distance between each curve, a tip velocity of 3 μm/s, a dwell time of 0.5 s and a set point of 1 nN was chosen. Prior to analysis, a baseline correction was performed for every force curve and the contact point was determined by visual inspection. A polynomial of the form 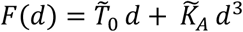 was fitted to the contact region to assess the apparent pretension 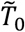 at low strain and the apparent area compressibilty modulus 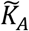 at large strain^20^. Baseline correction and analysis were performed with self-written MATLAB (The MathWorks, Natick, USA) scripts. Statistical analysis was performed with the Wilcoxon rank sum test.

For topography scans cells were fixed with 4 % formalin (Fluka, Buchs, Switzerland) in PBS for 20 min in order to obtain better lateral resolution. Scans were performed in contact mode with the same AFM and cantilevers and a set point force of 1.5 nN. Gains were adjusted so that no ringing was observed. A scan rate of 1 Hz or 0.5 Hz was chosen. For visualization, a line offset correction was performed for each scan line individually and a plane was subtracted from the image to correct for tilt in the sample.

### Electrical cell impedance sensing

A Zθ set-up from Applied Biophysics (Troy, USA) equipped with 8W1E electrode arrays (8W1E PET ECIS culture ware, Ibidi, Munich, Germany) was used and the measurements were carried out under cell culture conditions (32 °C and 5 % CO_2_). The electrodes were coated with collagen in the same manner as for the cell culture on petri dishes. With 400 μl FAD medium inserted per well, a baseline of the impedance was recorded for 1 h. Then, the medium was removed and 75,000 cells suspended in 400 μl FAD medium were inserted in every well. A change to FAD^+^ (high calcium) medium was performed roughly 30 h after seeding. One well was left without cells and one well per cell population was devoid of calcium during the whole measurement as a control. Measurements were performed at 11 different frequencies (ranging from 0.0625 kHz to 64 kHz) giving access to a complete impedance spectrum. For visualization of time traces at 1 kHz the real part of the impedance was baseline corrected by the values of only medium-filled electrodes. Complete spectra were normalized frequency-wise to electrodes filled with medium only. Fluctuation analysis was performed by calculating the power spectrum of the de-trended impedance signal of each well individually at steady-state conditions (*t* > 120 h) and fitting a slope to it.

## Results

### Inefficient wound closure and altered single cell mechanics in Kty^−/−^ cells

Keratinocytes are major building blocks of the epidermis and hence pivotal for epidermal wound healing after injuries. However, the role of the keratin network at the single cell level in this process is not entirely understood. To address this role in a quantitative and reproducible fashion, we used mechanically manipulated epithelial cell sheets by generating a single-cell wound in an otherwise intact monolayer.

Confluent cell sheets of WT and Kty^−/−^ cells were created by culturing the cells for 48 h in high calcium medium, before cortices of single cells were scratched with a micropipette. The treatment led to a void in the cell layer referred to as a single-cell wound (Figure 1 a). Successful treatment was inferred from the strongly reduced actin signal in harmed cells, indicating a loss of actin structures due to mechanical destruction of the filaments (Figure 1 b). In WT cells, the keratin network was still visible with merely a hole observed at the spot where the micropipette was inserted demonstrating that the keratin network was able to bear and endure the harsh mechanical treatment while the actin cytoskeleton was irreversibly compromised.

**Figure 1:**
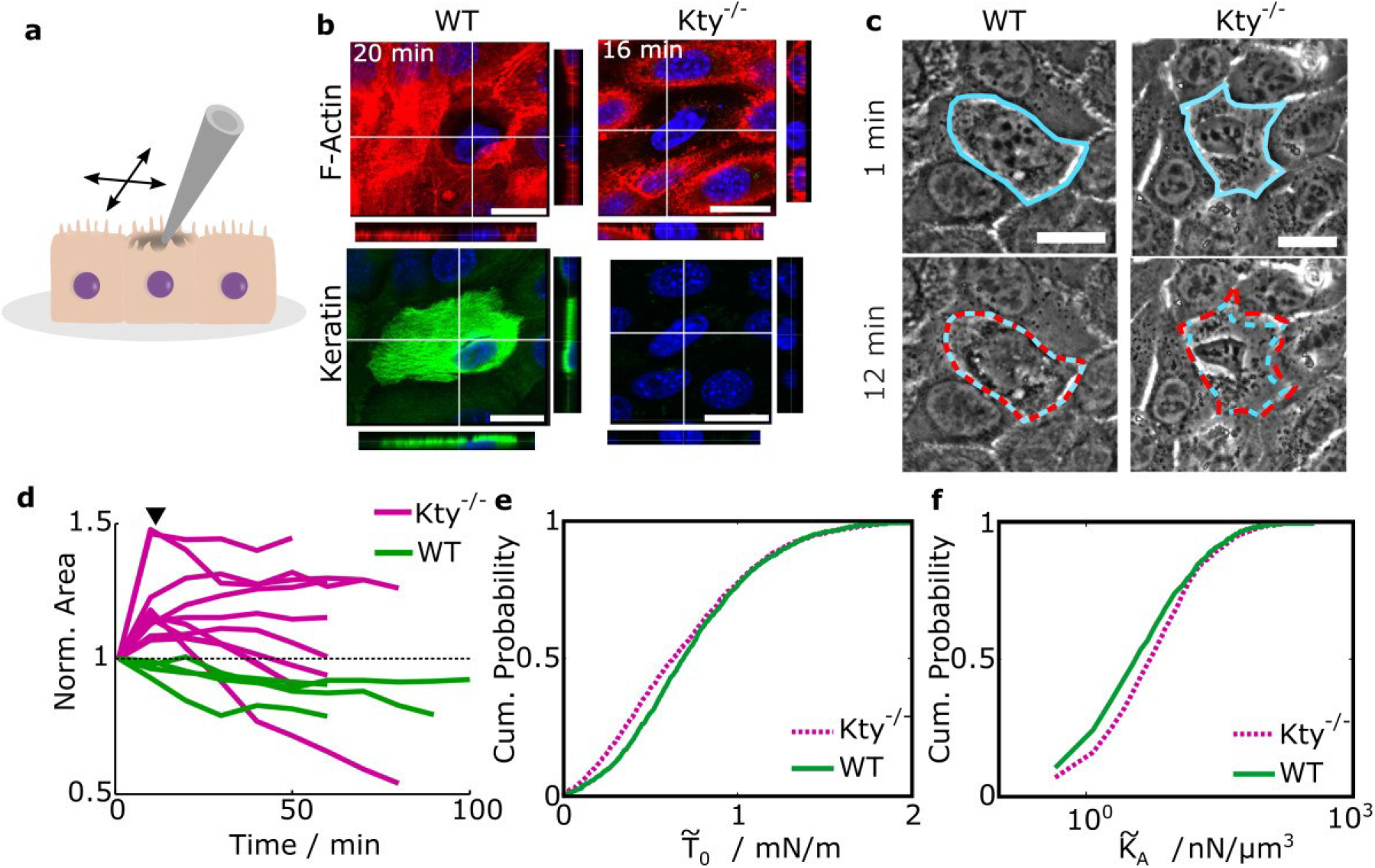
Inefficient wound closure in Kty^−/−^ cells: (a) A single cell in a confluent layer was scratched apically with a micropipette, leading to a single-cell wound. (b) Fluorescence micrographs including xz and yz sections of actin (red) and keratin 14 (green) in the wounded cell after micropipette-mediated wounding. (c) Phase contrast images of the indicated cell 1 min after wounding (blue contour) and 12 min after wounding (red contour). (d) Individual cell area changes of defect cells over time for WT (green) and Kty^−/−^ cells (magenta). Normalization was done with respect to the area prior to wounding. Arrowhead marks time point of 12 min which is visualized in c. (e) Cumulative histogram of the apparent pretension 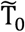 capturing the resistance to external force at low indentation depth (p<0.001) for homeostatic WT cells (green, solid line) and homeostatic Kty^−/−^ cells (magenta, dashed line). (f) Cumulative histogram of the apparent area compressibility modulus 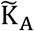 probing larger indentation depths (p<0.001) for the same cells. Number of analyzed force curves n = 2441 for Kty^−/−^ and n = 2563 for WT cells which corresponds to N = 5 individual experiments/force maps. Scale bars: 20 μm.

After wounding, defective WT cells retained their shape (Figure 1 c, Video SI1). In contrast, the cell body of Kty^−/−^ cells was ripped open and cytoplasmic content was pulled outwards (Video SI2). Detailed size analysis of the wounded region revealed an increase in size in Kty^−/−^ cells within the first 15 minutes after wounding (Figure 1 d). In contrast, WT cells instantly started wound closure by slowly decreasing the wounded cell area, independent of the size of the cell (Figure SI1). This pointed to a different mechanical relaxation process occurring in Kty^−/−^ cell sheets after wounding compared to WT cell layers, suggesting that the passive action of the keratin network might limit the immediate stress release.

Concerning single-cell mechanics of homeostatic cells, cells lacking the entire keratin network were found to be softer compared to WT cells^8–10^. We applied the previously introduced tension model to AFM force indentation experiments as it was also done for single-cell wound experiments^20^ and found that the apparent pretension dominated by cortical tension was decreased in Kty^−/−^ cells (Figure 1 e). In contrast, the second parameter obtained from the tension model, namely the apparent area compressibility modulus typically associated with excess area and the elastic modulus of the cell cortex, was increased in Kty^−/−^ cells compared to WT cells (Figure 1 f). A possible explanation of this increase in Kty^−/−^ cells will be given below during analysis of the cell’s cytoskeletal organization.

### Loss of mechanical coupling in Kty^−/−^ epidermal sheets

To investigate whether and to which extent keratins were involved in the response of neighboring cells to mechanical loading of a single cell by micropipette wounding, WT and Kty^−/−^ cells were analyzed by video microscopy during single-cell wounding. The videos revealed that only movements of the actually scratched cell were visible in Kty^−/−^ cells (Video SI3), whereas in WT cells, movements of entire cells and cell junctions one cell width away from the wounded cell were detected (Figure 2 a, Video SI4), indicating a mechanical coupling in the cell layer during the stimulus. For Kty^−/−^ cells, no movement was detected at junctions one cell separated from the stimulus (Figure 2 b, Video SI3), suggesting lack of keratins had compromised the mechanical coupling within the single cell layer.

**Figure 2:**
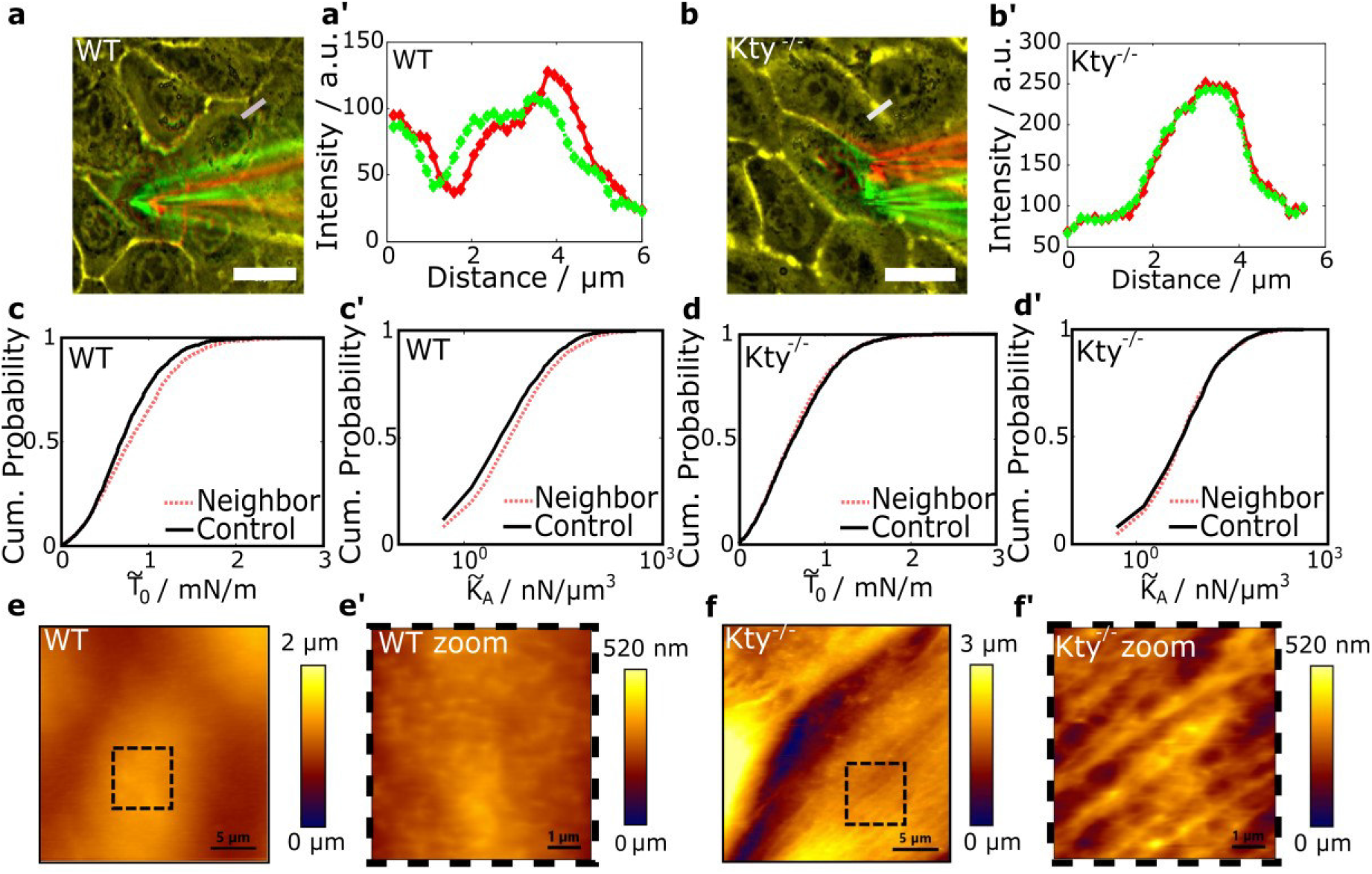
Impaired mechanical coupling in the absence of keratins: (a) Merged image of colored (red, green) phase contrast images taken during wounding of WT cells. Yellow color indicates exact overlay of images, red colored image was taken before micropipette movement, green after movement. (a’) Line profiles of the two images at a position of a cell junction one cell away from the defect (grey bar in a) indicating junction movement during wounding. (b-b’) Same analysis for Kty^−/−^ cells during wounding. (c) Cumulative histogram of the apparent pretension 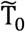 and (c’) apparent area compressibility modulus 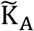 of WT cells neighboring a defect (red, dashed) and control cells in an intact layer (black, solid) (both p<0.001; n = 2659 (Neighbor) and n = 2563 (Control)). (d-d’) Cumulative histograms of the same mechanical parameters for Kty^−/−^ cells (p=0.112 and p=0.288, respectively; n = 2225 (Neighbor) and n = 2441 (Control)). (e) AFM height scan of WT cells revealing homogeneous junctions. (e’) Zoom into marked area in e’ showing homogeneous cortex structures. (f) AFM height scan of Kty^−/−^ cells revealing fragmented junctions. (f’) Zoom into marked area in f’ showing fibrous structures. Scale bars: (a) and (b): 20 μm, (e) and (f): 5 μm, (e’) and (f’): 1 μm.

Moreover, we found that mechanical responses to single-cell wounding in cells neighboring the defect differ depending on the presence or absence of keratins. In close analogy to previous findings in keratin-expressing MDCK II cells, neighboring WT keratinocytes showed larger mechanical resistance mirrored in both parameters, the apparent pretension and the apparent area compressibility modulus (Figure 2 c)^20^. Kty^−/−^ cells, however, showed the same single-cell mechanics in control cells surrounded by an intact layer and defect neighboring cells (Figure 2 d). This indicated that keratin-deficient cells do not respond to mechanical wounding of neighboring cells. As a consequence, wound response, single-cell mechanics, and mechanical coupling were all altered in keratinocyte cell sheets lacking a keratin network. Next, we turned to the identification of cell components mediating this behavior. Given that the observed loss of coupling between cells indicated changes in junctional complexes in cell sheets, we reasoned that cortical tension generation which depends on the integrity of intercellular junctions, might be compromised in the absence of keratins^24^.

Indeed, atomic force microscopy imaging confirmed the impairment of the junction zone (Figure 2 e and 2 f) in keratin-deficient cells. In WT cells, the topography of the junction area was continuous in height while in Kty^−/−^ cells distinct elevated structures occurred at the position where neighboring cells were in contact with each other. In addition, gaps between neighboring cell membranes were found. Furthermore, thick fibrous structures across the cell body were visualized in Kty^−/−^ cells (Figure 2 f’). These structures were not detected in WT cells, in which a homogeneous height profile across the cell body was found (Figure 2 e’). We hypothesize that these thick bundles under the cell’s surface might also explain the large resistance found for Kty^−/−^ cells during AFM indentation experiments at larger cell penetration depths.

This prompted us to analyze differences in the composition of junctions and the associated actomyosin structures.

### Altered acto-myosin organization in the absence of keratins

In line with the AFM data, F-actin staining showed that junctional actin fibers were significantly reduced in the absence of keratins compared to controls. Conversely, the number of stress fibers spanning the cytoplasm was significantly increased in Kty^−/−^ keratinocytes compared to WT cells (Figure 3 a-d). In contrast, microtubules appeared to be unaffected in Kty^−/−^ cells (Figure SI 2 c). In general, reorganization of the actin cytoskeleton together with tightly controlled myosin-II activity produces mechanical forces that drive junction assembly, maintenance, and remodeling^25^. To investigate whether in addition to impaired reorganization of the junctional actin cortex, loss of keratins also caused changes of myosin-II activity, WT and Kty^−/−^ cells were stained for total (MLC2) and active, phospho-myosin light chain (pMLC). In WT cells, total and active myosin staining overlapped with the junctional actin cortex and was homogenously distributed around cell-cell contacts (Figure 3 a’-a’’’ and c’-c’’’). In contrast, in Kty^−/−^ cells active myosin light chain was concentrated in a punctate pattern along cell-cell borders and overlapped to a great extent with intracellular actin stress fibers (Figure 3 b’-b’’’ and d’-d’’’). Western blotting for active myosin light chain and total myosin light chain revealed a significant decrease of myosin-II activity in the absence of keratins (Figure 3 e-e’). This suggests that in addition to altered actin organization, overall actomyosin contractility is decreased in the absence of keratins.

**Figure 3:**
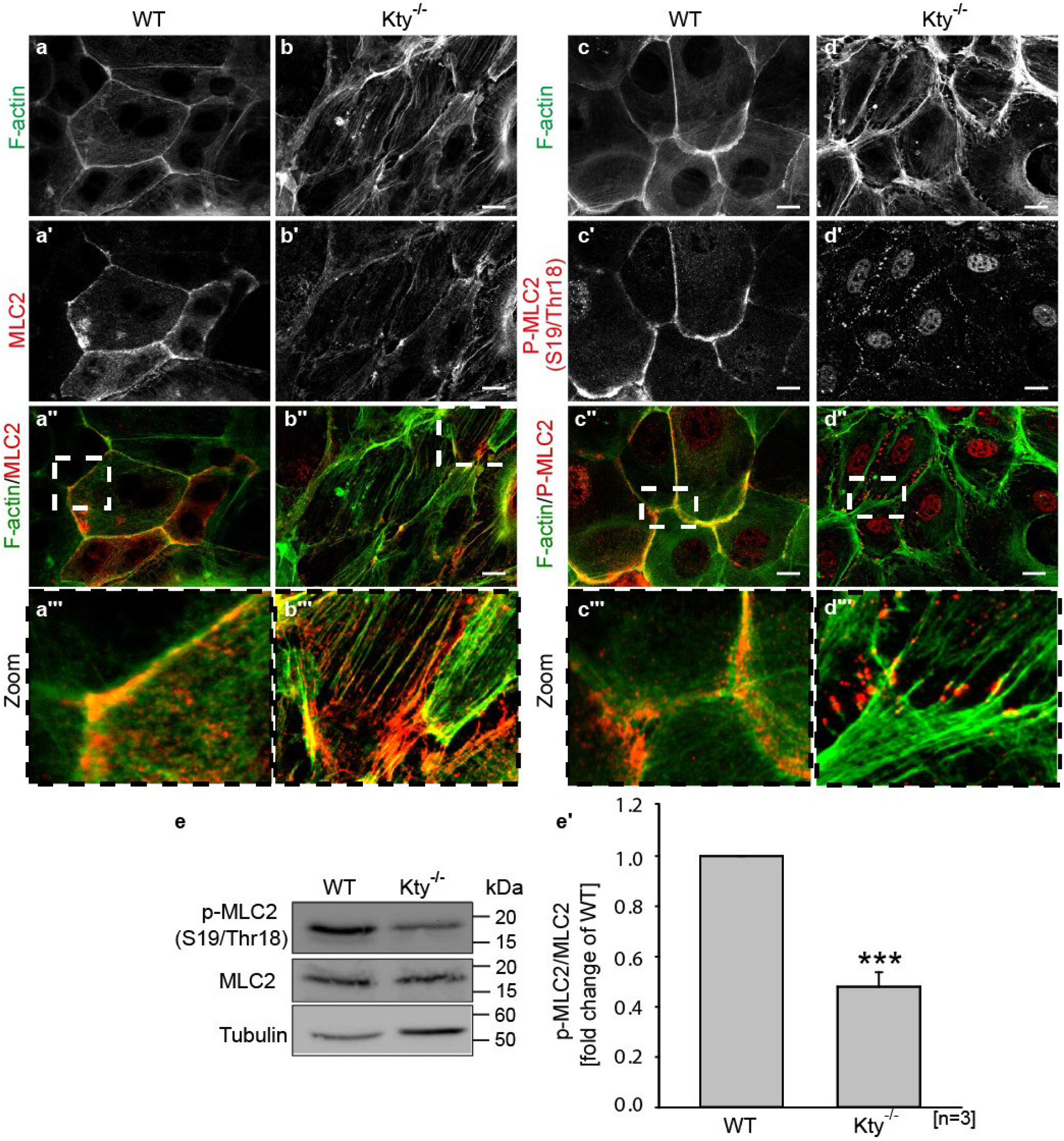
Altered actomyosin organisation in Kty^−/−^ cells: (a-d’’’) Immunostaining for F-actin (Phalloidin-Alexa 488), MLC2 and phospho-myosin light chain (pMLC) in WT and Kty^−/−^ cells 48 h after Ca^2+^-switch. Scale bars: 10 μm. (e) WB of p-MLC and MLC (e’) Quantification of P-MLC-levels relative to total MLC2 (mean+/−SEM, n=3).

### Impaired formation of cell-cell junctions in Kty^−/−^ cells

The impaired mechanical coupling of keratin-deficient keratinocytes and the reorganization of the actin cytoskeleton prompted the question whether in addition to compromised desmosomes in the absence of keratins^10,11^, actin-linked adherens and tight junctions were affected. Immunofluorescence staining of E-cadherin (Ecad), α-catenin, and p120-catenin showed that Kty^−/−^ cells formed only an immature, “zipper-like” appearance of adherens junctions (Figure 4 a-c’), whereas in WT cells, adherens junction proteins were localized as a straight line tightly connected to the plasma membrane (Figure 4 a-c’). This altered organization was accompanied by slightly reduced levels of adherens junctions proteins in Kty^−/−^ cells 48 h following the calcium-switch (Figure 4 d-d’).

**Figure 4:**
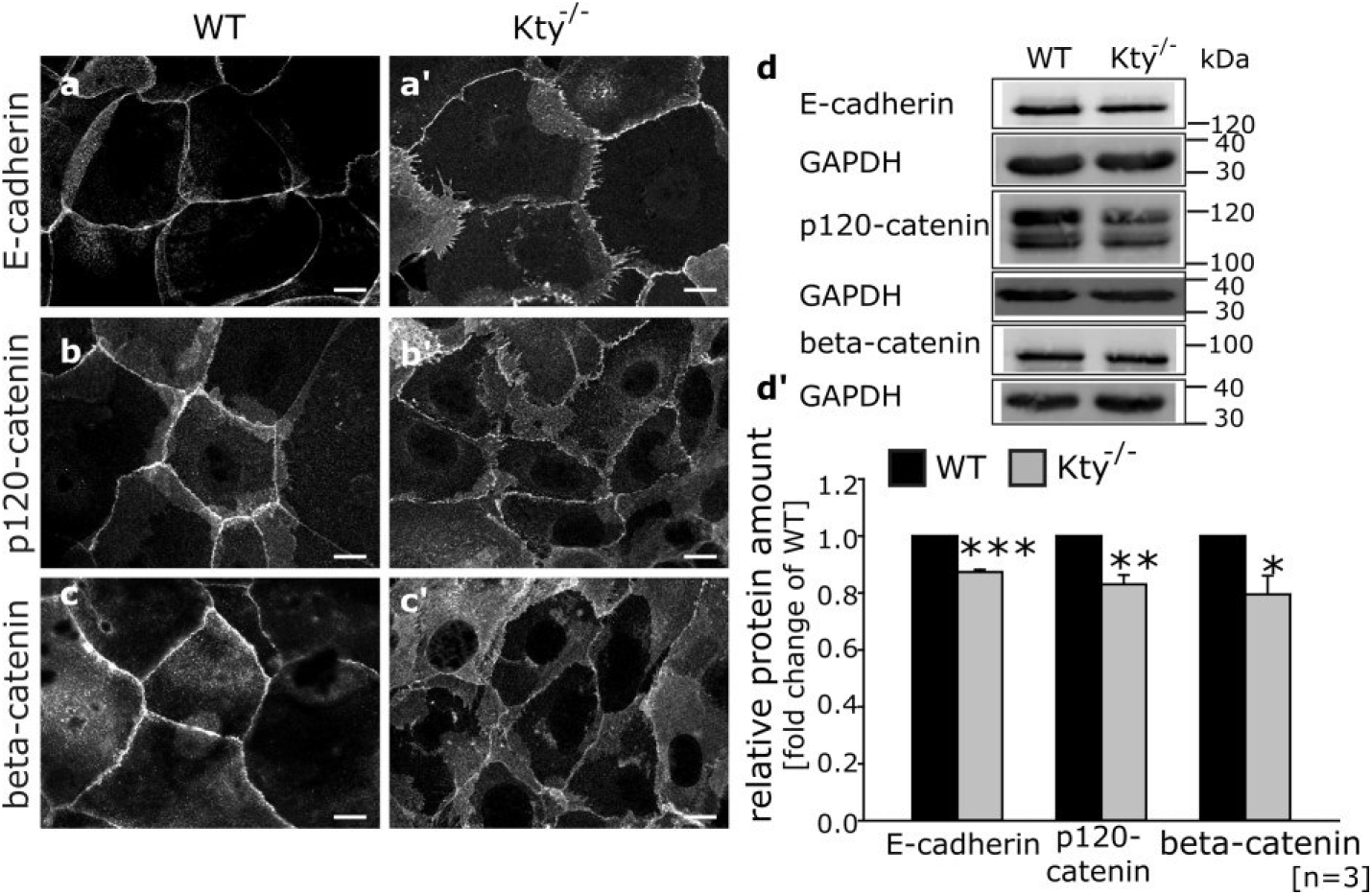
Impaired formation of adherens junctions: (a-c’) Immunostaining for E-cadherin, p120-catenin, and beta-catenin in WT and Kty^−/−^ cells 48 h after Ca^2+^-switch. Scale bars: 10 μm. (d) WB of total protein lysates for E-cadherin, p120-catenin, beta-catenin, and Tubulin. (d’) Quantification of protein levels relative to tubulin as loading control (mean+/−SEM, n=4).

In addition to adherens junctions, tight junction organization was severely impaired in the absence of keratins. Although occludin is still localized to cell-cell borders in Kty^−/−^ cells, its distribution appeared very inhomogeneous in comparison to WT cells (Figure 5 a-d). Moreover, claudin 1 and 4 were almost completely absent from cell-cell junctions in keratin-free keratinocytes (Figure 5 a‘-d‘‘). Western blot (WB) analysis of total protein lysates revealed a significant decrease of occludin, claudin 1, and claudin 4 protein levels in keratin-free keratinocytes (Figure 5 e-e‘). The impaired organization of adherens and tight junctions in keratin-deficient cells indicated that besides actin organization, also the formation of cell-cell contacts was disturbed in the absence of keratins underlining the findings described above from AFM topography scans.

**Figure 5:**
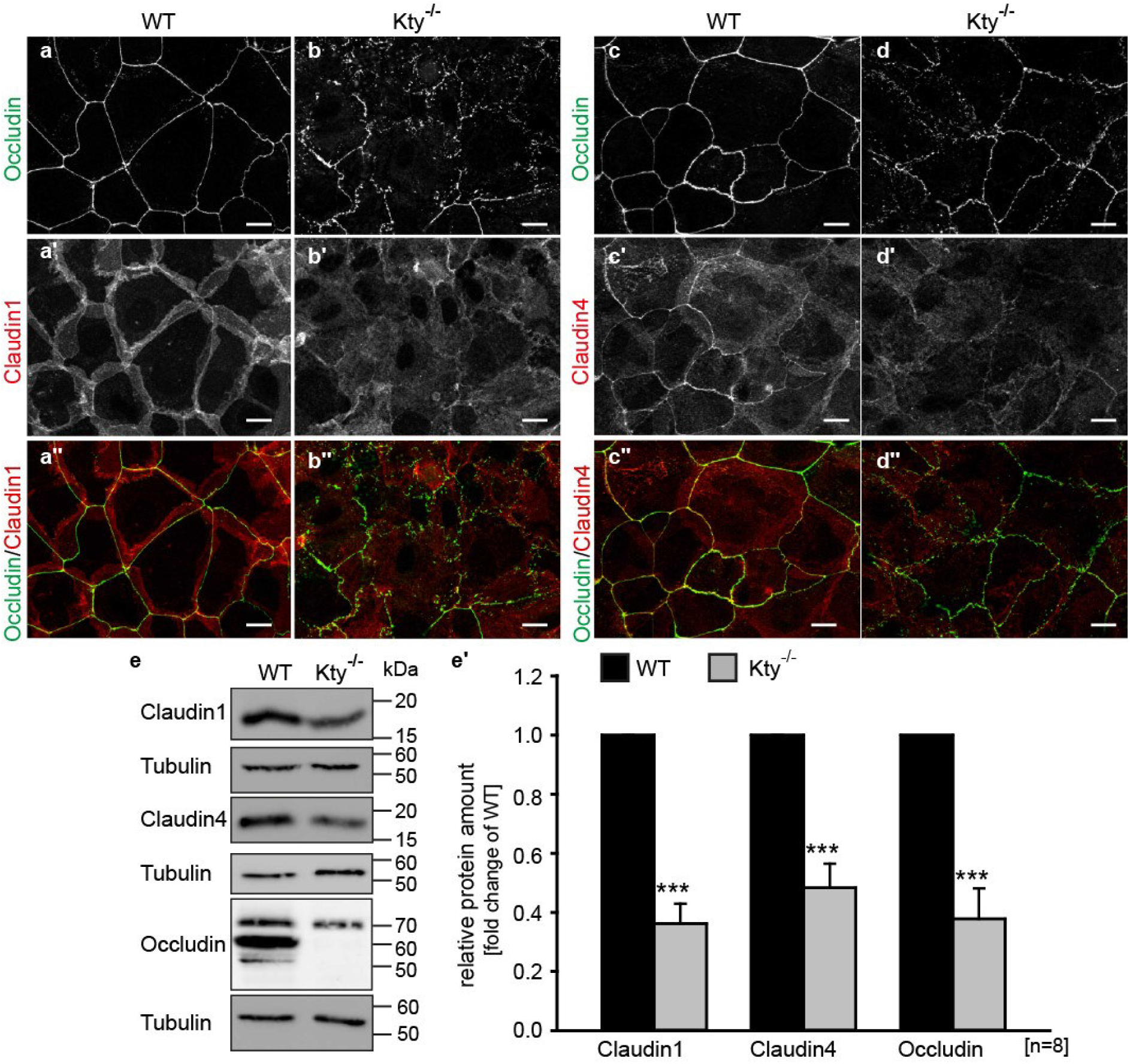
Altered organization of tight junctions in keratin-free keratinocytes: (a-d’’) Immunostaining for occludin, claudin 1, and claudin 4 in WT and Kty^−/−^ cells 48 h after Ca^2+-^ switch. Scale bars: 10 μm. (e) WB of total protein lysates for occludin, claudin 1, claudin 4, and α-tubulin. (e’) Quantification of protein levels relative to α-tubulin as loading control (mean+/−SEM, n=8).

### Impaired barrier function in Kty^−/−^ epidermal cell sheets

The extra-membranous distribution of occludin and claudins 1 and 4 suggested an impaired tight junction barrier^26^. In order to substantiate a dysfunctional tight junction barrier, electrical cell-substrate impedance sensing was performed, which measures the sealing efficiency of tight junctions by applying an alternating current and monitoring the resulting complex impedance Z due to current flow through the paracellular space or the membrane. The real part of the impedance at an intermediate frequency regime of 200 Hz < f < 5 kHz, indicative for sealing of the paracellular space by junctions^27^, was monitored over time. Three different phases were identified (Figure 6 a). Before calcium addition, both WT and Kty^−/−^ cells generated the same low impedance (Phase I named P I) indicative of premature, leaky cell-cell contacts. Residual impedance originated from cells adhering to the gold electrodes and thereby limiting the ionic conductance. Instantaneously after calcium addition, a pronounced barrier against ion flux formed in WT cells as inferred from an increased impedance in P II. At the same time, Kty^−/−^ cells showed impedance levels matching that of the ‘low calcium’ control cells, indicating that no barrier had formed and junction sealing was incomplete for these cells. Only at time frames when WT cells already reached a stable barrier (named P III), Kty^−/−^ cells also started to establish a detectable barrier. However, even upon impedance saturation, Kty^−/−^ cells still showed lower values in comparison WT cells. Differences in impedance increase upon layer formation were seen for all intermediate frequencies (Figure 6 b). As the high frequency regime is mostly dominated by surface coverage and not influenced by junction formation^27^, differences between WT and Kty^−/−^ vanished there, indicating that indeed junction formation caused the differences in impedance levels between WT and Kty^−/−^ cells.

**Figure 6:**
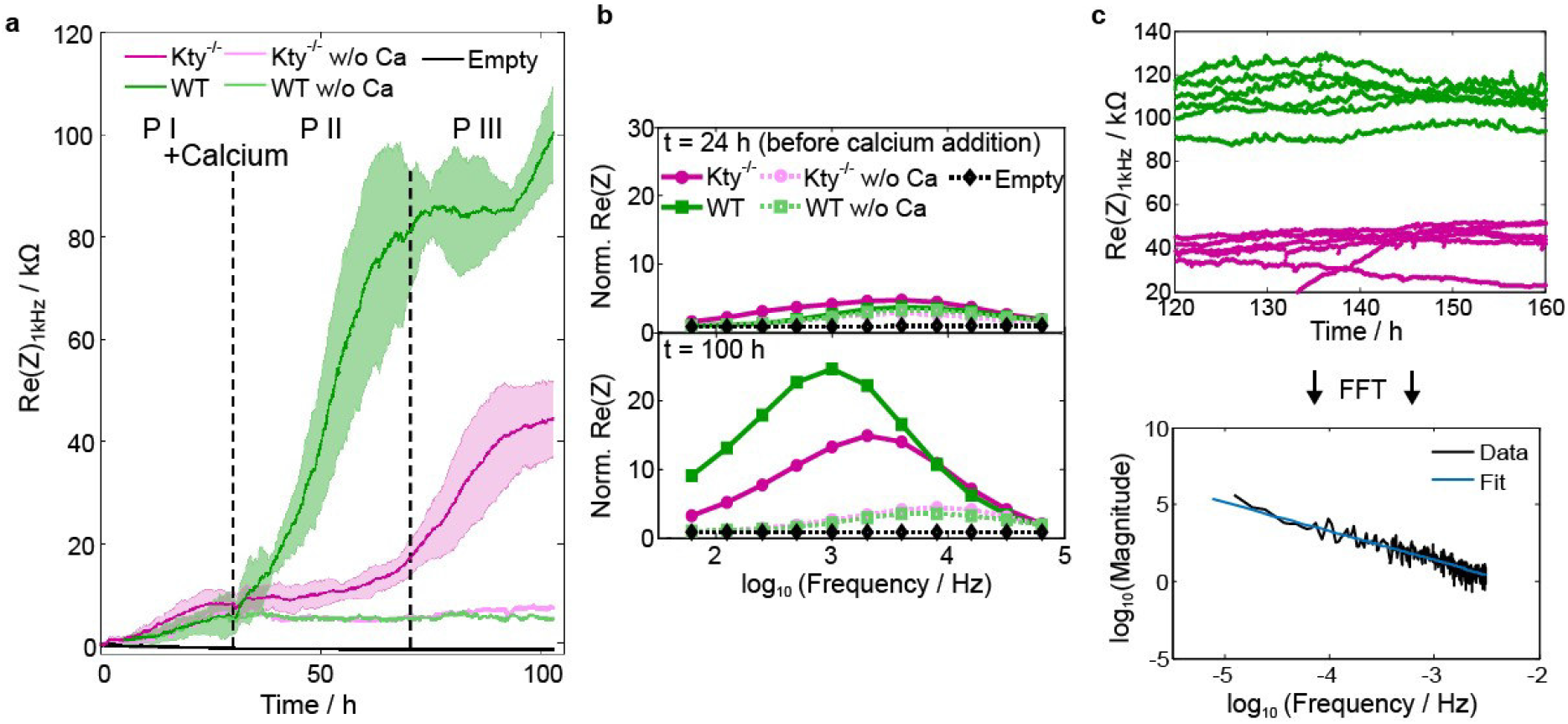
Barrier formation is compromised in Kty^−/−^ cells: (a) Real part of the complex impedance Z recorded with electrical cell-substrate impedance sensing at 1 kHz as an indicator for junction functionality followed over time for WT (green, N=4), Kty^−/−^ (magenta, N=4), ‘low calcium’ controls (light green, light magenta, respectively) and empty electrodes (black). (b) Normalized impedance spectra at t = 24 h (before calcium addition) and after impedance leveling in Kty^−/−^ cells at t = 100 h. (c) For fluctuation analysis each individual signal is Fourier transformed (black) and the resulting power spectrum is fitted with a linear slope (blue).

Furthermore, impedance monitoring allows for fluctuation analysis of the cell-substrate distance and cell-cell contacts often referred to as micromotion. It was found that the spectral behavior of micromotion time series can be related to the viability of cells and their general biological activity. We therefore analyzed impedance fluctuations under steady-state conditions (*t* > 120 h) and found that the signal showed larger fluctuations for WT cells than for Kty^−/−^ cells (Figure 6 c). Not only is the variance larger but also the memory of impedance fluctuations is different. The resulting power law dependence ∝ *f*^−*β*^ in the power spectrum with *β* being generally between 0.6 and 2.5 is indicative for the presence of a long time memory effect in the electrical signal^28^; it basically represents the Hurst coefficient *H* of the noise (*H* = (*β*-1)/2). A value of *H* in the range of 0.5–1 indicates a time series with long-term positive autocorrelation. This means that a high value in the impedance time series will probably be followed by another high value and that the expectation for data in the following time points will also be that higher impedance values occur. Values larger than *β* > 2 are indicative of fractional Brownian motion. Indeed, the slope in the power spectrum of WT cells revealed a value of *β*_WT_=(2.02±0.18) (*N*=12). This magnitude in slope indicates that the fluctuations underlie a long-term memory effect and active noise might be present. In Kty^−/−^ cells, the slope decreased to *β*_Kty−/−_= (1.75±0.18) (*N*=12). In earlier studies values of around 2 and larger were found for living cells of different cell lines while *β* for the same cells fixed with glutaraldehyde were found to be around values of less than 1^28^. This indicates that junction regulation in WT cells depends on active contributions while Kty^−/−^ cells show substantially less activity.

Thus, the functional analysis of tight junctions revealed an impaired barrier function in Kty^−/−^ cells and showed that long-term memory effects of junctional conductivity were higher in the presence of keratins.

### Re-expression of K5/K14 restores wound closure and cell adhesion but not the full barrier functionality

Thus far, our experiments have uncovered that the complete loss of a keratin network has crucial implications for wound closure, mechanical coupling and barrier formation within a keratinocyte cell sheet. To examine to which extent single cell wound recovery, cell adhesion and tight junction barrier depended on keratins, K14 were re-expressed in Kty^−/−^ cells (subsequently referred to as K14 cells), which stabilizes the endogenous K5 and led to the formation of a K5/14 keratin network^10,29^. Previously, we showed that rescue cells express ~38% of total keratin levels in WT controls^10,29^.

Concerning wound closure, the wound size progression after defect-induction mirrored the WT behavior, i.e., the generated gap area remained constant or was even slowly decreasing over time. It progressed without intermittent enlargement of the wound area, which was, however, typical of Kty^−/−^ cells (Figure 7 a, Video SI5). In addition, mechano-coupling between freshly wounded and neighboring cells was restored, as the latter cells showed movements similar to those found for WT cells (Video SI6). In further support, line profile analysis revealed a movement of the junction position in cells which were not directly affected by the micropipette movement (Figure 7 b). However, we noted that despite the presence of K5/K14, neighboring cells still failed to show an increased stiffness in response to wounding during the closure process (Figure SI2 a-a’), which was found for WT cells.

Immunostainings of adherens and tight junction proteins showed a homogenous distribution at cell-cell junctions, similar to WT cells (Figure 7c-e’’, SI2 d). This distribution of cell-cell junctions was mirrored in AFM topography scans, which revealed homogenous apical structures and junction appearance in K14 cell sheets (Figure SI2 b), resembling those in WT cells. Western blotting for tight junction components showed a partial increase of total levels of claudin 1, claudin 4, and occludin after re-expression of K5/K14 (SI2 e). Furthermore, the cytoplasmic arrays of actin stress fibers were reorganized into peripheral actin rings in rescue cells, similar to those observed in WT controls (Figure 7 f-f’’). As before, microtubule staining appeared unaffected by keratin expression in all three cell-lines (Figure SI 2 c-c’’’).

**Figure 7:**
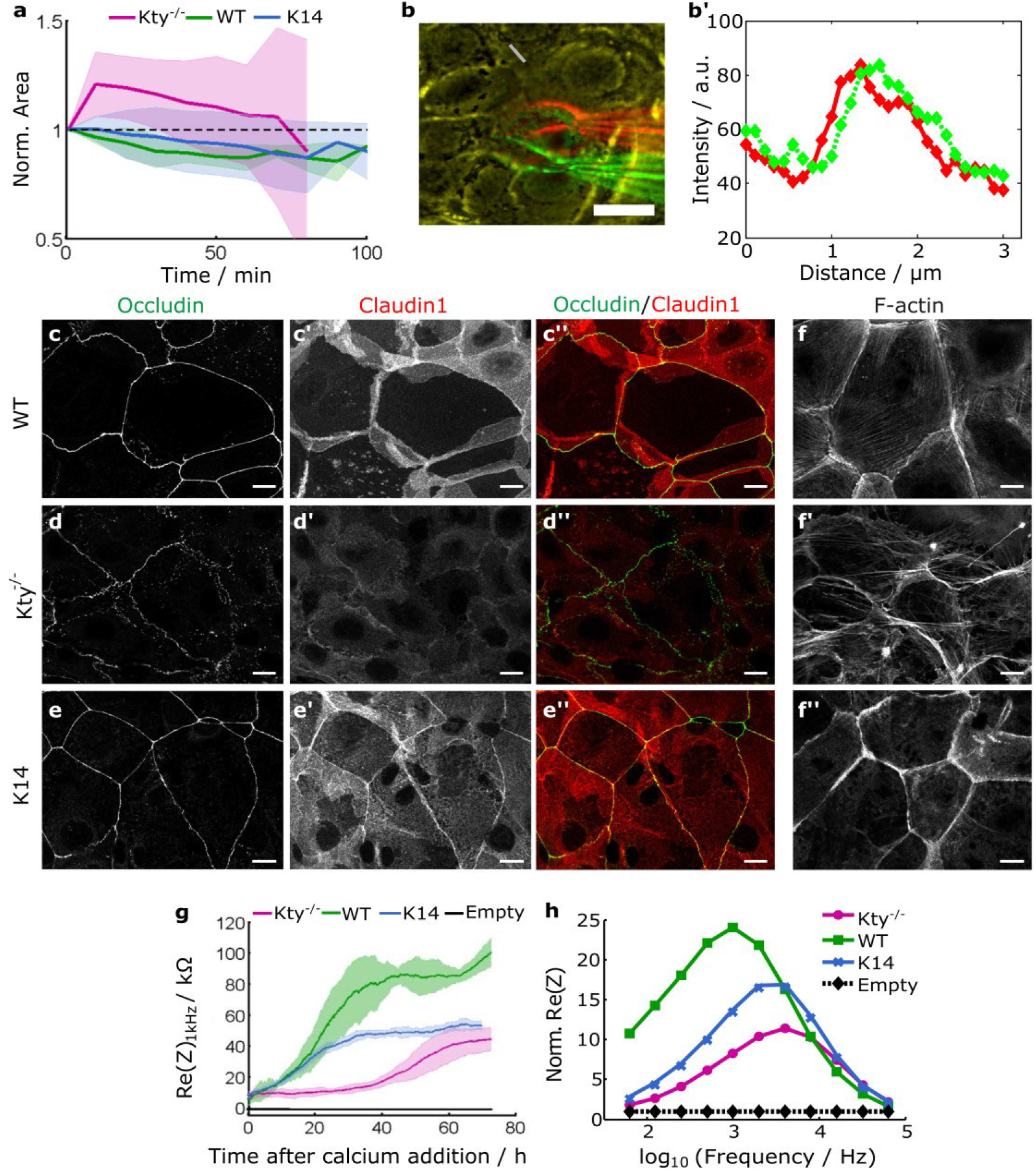
Re-expression of K14 partially restores TJ organization and barrier formation: (a) Averaged cell area changes of defect cells over time for WT (green, N = 4), Kty^−/−^ cells (magenta, N = 9) and K14 cells (blue, N = 7) including standard deviations (shaded areas). Normalization was computed with respect to the individual area shortly before wounding. (b) Merge of colored phase contrast images taken during wounding in K14 cells. Yellow color indicating exact overlay of images, red image was taken before micropipette movement, green after movement. Scale bar: 20 μm. (b’) Line profiles of the two images at a position of a cell junction one cell away from the defect (grey bar in b) indicating displacement of junctions during wounding. (c-e’’) Immunostaining for occludin and claudin 1 in WT, Kty^−/−^, and K14 rescue cells 48 h after Ca^2+^-switch. Scale bars: 10 μm. (f-f’’) Immunostaining for F-actin (Phalloidin-Alexa 488) in WT, Kty^−/−^, and K14 rescue cells 48 h after Ca2+-switch. Scale bars: 10 μm (g) Real part of the impedance Z recorded at 1 kHz as a representative indicator for junction functionality followed over time for WT (green, N = 4), Kty^−/−^ (magenta, N = 4), K14 rescue cells (blue, N = 4), and the empty electrode (black). (h) Normalized impedance spectra (real part) 48 h after calcium addition.

Electrical barrier measurements using electrical cell-substrate impedance (ECIS) revealed an instantaneous barrier formation in K14 cells after calcium addition, similar to WT cells (Figure 7 g). However, the impedance at 1 kHz at steady-state remained between that of WT and Kty^−/−^ cells. Recordings of complete spectra showed that rescue cells display impedance values between those of WT and Kty^−/−^ cells 48 h after calcium addition (Figure 7 h). Furthermore, fluctuation analysis proved that K14 cells showed the same level of active noise in impedance measurements as WT cells (with β_K14_ = (2.07±0.20) (N = 12) and β_WT_ = (2.02±0.18), while knock-out cells arrived merely at β_Kty−/−_ = (1.75±0.18)). Consequently, the ability to form a functional barrier was partially restored by re-expression of K5 and K14 filaments.

In conclusion, the functionality of junctions and of the cytoskeleton was partially rescued by the re-expression of K5/K14 in Kty^−/−^ keratinocytes. The rescued cells were not fully capable of reestablishing the original barrier and to undergo wound-induced stiffening as WT cell do. This indicates that relative keratin abundance is a crucial determinant of micromechanical cell properties and tight junction functionality^10,29^.

## Discussion

This study highlights the importance of an intact keratin network for mechanical integrity and tight junction-dependent barrier functionality in keratinocyte cell sheets. Moreover, it demonstrates that lack of keratins affects the function of adherens and tight junctions and thereby compromises the diffusion barrier (Figure 4,5). The absence of keratins caused extensive rearrangement of the cells’ contractile elements, leading to the appearance of thick cytoplasmic stress fibers (Figure 2 e-f’, Figure 3). We posit that this change in the contractility pattern of individual cells can explain most of the findings presented here.

It is known that adherens junctions are regulated by the contractility of the actin cytoskeleton^30^, as force positively influences adherens junction size^31^. Also, it is known that actin regulates the barrier efficiency in epithelial cells by means of contractile forces applied at tight junctions^32^ which can extend to tight junction positioning within a tissue^33^. Furthermore, establishment of adherens junctions and tight junctions is locally coupled^34^. If force applied intracellularly at junctions is very localized and limited to distinct spots, irregular positioning of junctions occurs, explaining the loss of barrier functionality as seen in ECIS experiments (Figure 6). Moreover, loss of memory in the impedance signal points to a less coordinated regulation of junctions over time, which can originate from impaired actomyosin contractility.

Indeed, the analyzed impedance fluctuations obtained from the frequency regime where TJ prevail (here 1 kHz was chosen) indicate that concerted action exists only in cells with an intact keratin network.

A tight regulation of contractile and load-bearing elements is as crucial for single cells as for collective cell mechanics. According to the tensegrity model, a composite network of filaments with different mechanical properties build up a flexible yet mechanically stable three-dimensional architecture inside individual cells^35^.

We therefore suggest that with the rearrangement of contractile actomyosin inside the cell, cells try to compensate for the loss of keratin filaments to maintain mechanical homeostasis: keratin filaments prevent the cells from rupture upon large strain, which actin cannot compensate due to different filament mechanics^36^. This can also be seen during wounding, where mechanical load in WT cells is transmitted to neighboring cells through junctions further away from the actual micropipette movement (Video SI 3-4, Figure 2 a’-b’’). In Kty^−/−^ cells this mechanism is suppressed and only individual cells have to cope with local mechanical disturbance (Figure 1 c,d). As these cells are able to spread, grow and form junctions, some kind of mechanical homeostasis necessary for survival remains established in keratin-deficient cells. However, if contractility in one cell is externally disturbed as after wounding due to micropipette movement resulting in the loss of actin filaments, Kty^−/−^ are not capable to rely on established cytoskeletal elements for compensation and wound closure. This resulted in an increase in wound area and eventually rupture of the cell layer.

In general, it was demonstrated that epithelial cells in a homogeneous, homeostatic layer exist in a pre-tensed state^37^ and upon large scale wounding, the tissue locally relaxes as indicated by an initial increase in wound size^38^. However, upon single-cell wounding, neither MDCK II cells nor WT keratinocytes showed an increase in the gap size directly after wounding, as found for Kty^−/−^ cells. Therefore, we hypothesize that Kty^−/−^ cells are under a larger uncompensated pretension, which is released upon micropipette wounding and loss of cortical contractile fibers. The mechanism can be nicely illustrated with a spring model accounting for actomyosin contractility (Figure 8): In Kty^−/−^ cells, tensed mechanical springs are arranged in series, where a single spring symbolizes a single contractile bundle connected to a neighboring spring at spots where cell-cell junctions are perpendicular to the cell boundary. If all springs inside one cell fail instantaneously, the springs in neighboring cells recoil to a new resting length and the area of the wound increases consequently. In WT cells, however, a well-connected homogeneously contracted cortex exists which stabilizes the layer even upon loss of all contractile elements inside one cell. Intermediate filaments might act as shock absorbers in this process, i.e., dissipating the energy effectively.

**Figure 8:**
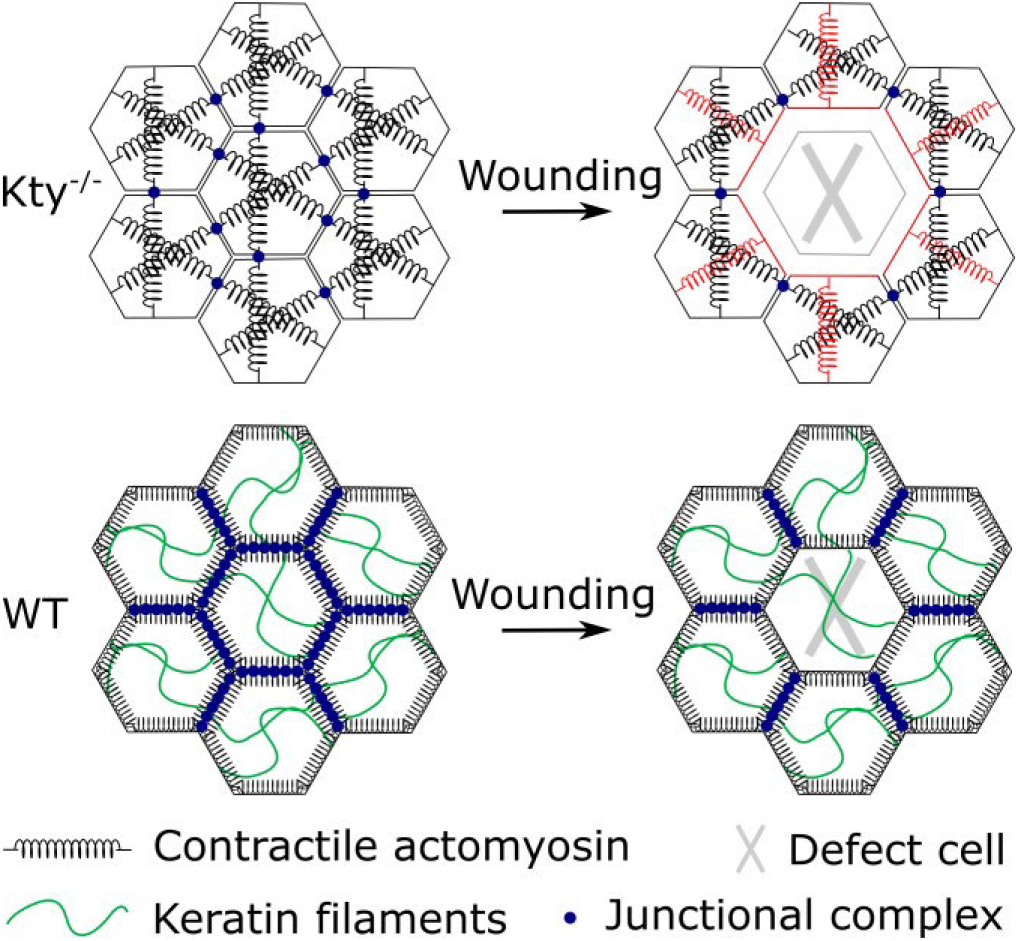
Schematic model of the contractile pattern inside Kty^−/−^ (top) and WT cells (bottom): Actomyosin in Kty^−/−^ cells is bundled into thick fibers spanning the cell (black springs). Only at distinct spots, namely the respective end of the fibers, cells are connected via cell-cell junctions (blue dots). In WT cells, a contractile cortex exists, lining the cell. Keratin filaments (green) are distributed inside cells acting as shock absorbers. Cell-cell junctions are distributed all over the cell’s periphery. Upon wounding and concomitant loss of contractile filaments inside one cell (grey X), adjacent contractile filaments are directly affected. In surrounding Kty^−/−^ cells, forces are not balanced anymore so that contractile filaments shrink and pull cell borders (indicated by red springs and borders) while in WT cells forces are distributed more evenly and stable junction positions and resting lengths of the contractile filaments are possible.

Hence, in WT cells the junctions stay local upon wounding before a slow closure process starts. Here, the neighbors even strengthen their cortex which becomes manifest in the increased mechanical pretension found in adjacent cells (Figure 2 c’-d’’).

In K14 rescue cells, a mixture of both settings was observed (Figure 7). Junctions and cortex were more homogeneous than in keratin-deficient cells and micropipette movements were transmitted throughout the layer. Also, the cells exhibit a reduced unbalanced contractility compared to Kty^−/−^ cells as indicated by the wound size progression. However, tight junctions remained altered, as seen by ECIS measurements, which points to persisting discrepancies compared to WT cells. A possible explanation for the incomplete rescue of tight junction functionality is that the overall keratin abundance was reduced in rescue compared to WT cells^10,29^. Alternatively, keratin isoforms that might be required for the formation of an intact barrier might still be lacking in K14 cells^17^.

In summary, we identified a novel role of keratins in controlling tight junctions and actomyosin contractility with major implications for wound closure and barrier functionality. This highlights the importance of keratin IFs for tissue function and mechanical homeostasis in keratinocyte cell layers.

## Acknowledgements

The authors thank T. Oswald (University of Göttingen) for critical reading of the manuscript and experimental support and Angela Ruebeling (University of Göttingen) for cell service and technical assistance. We greatly acknowledge the funding by DFG: SPP 1782 to A. J. and T.M.M.

## Author contributions

S.K. and F.B. carried out experiments and analyzed data. All authors wrote the manuscript.

