## Supplementary Information for "An intact keratin network is crucial for mechanical integrity and barrier function in keratinocyte cell sheets"

### Supplementary Table 1 Antibodies

| **Primary antibodies** | **dilution IF/WB** | **host** | **source** |
|---|---|---|---|
| E-cadherin (DECMA-1) | 1:500 (IF) | rat | Sigma Aldrich |
| E-cadherin | 1:1.000 (WB) | rabbit | BD Transduction Laboratories |
| β-catenin (Clone 14) | 1:100/1:2.000 | rabbit | BD Transduction Laboratories |
| p120-catenin (Clone 98) | 1:100/1:1.000 | rabbit | BD Transduction Laboratories |
| α-Tubulin | 1:12.000 (WB) | mouse | Sigma Aldrich |
| Claudin-1 | 1:400/1:20.000 | rabbit | Gift Niessen Lab (Cologne) |
| Claudin-4 | 1:400/1:400 | rabbit | Thermo Scientific |
| Occludin (OC-3F10) | 1:250 /1:400 | mouse | Thermo Scientific |
| ZO1 | 1:250 (IF) | rabbit | Thermo Scientific |
| MLC2 | 1:100/1:1.000 | rabbit | Cell Signaling |
| P-MLC2 (T18/S19) | 1:100/1:1.000 | rabbit | Cell Signaling |
| **Secondary antibodies** | **dilution** | **host** | **source** |
| anti-mouse-, anti-rabbit-, anti-rat-, anti-guinea-pig-DL488, 549 | 1:800 | donkey | Dianova, Hamburg |
| anti-mouse-, anti-rabbit-, anti-rat-, anti-guinea-pig-DL649 | 1:100 | donkey | Dianova, Hamburg |
| anti-mouse-, anti-rabbit-HRP | 1:20.000 | donkey | Dianova, Hamburg |

### Figures

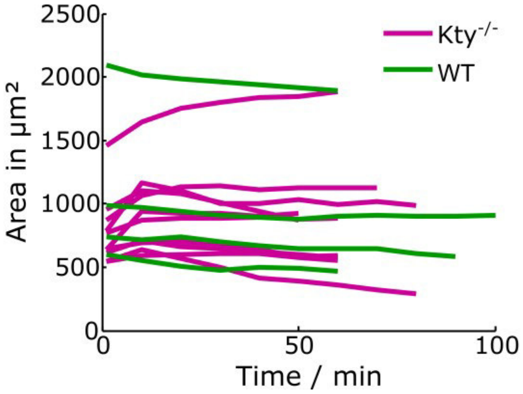


Figure SI1: Absolute values of wound area progression of individual cells over time in Kty$^{-/-}$ cells (magenta) and WT cells (green).

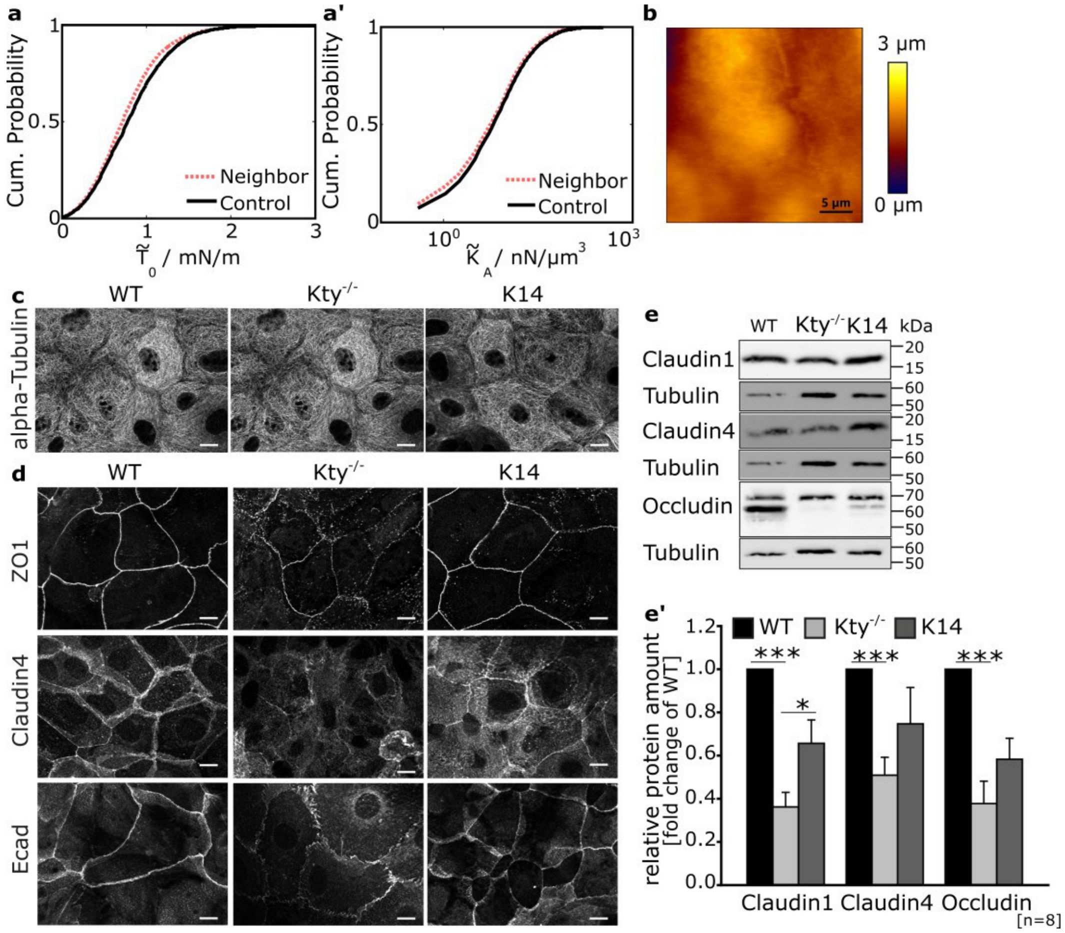


Figure SI2: (a) AFM-based mechanical analysis showing the cumulative distribution of the apparent pretension $\tilde{T}_0$ and (a') the apparent area compressibilty modulus $\tilde{K}_A$ of K14 cells neighboring a defect (red, dashed) and control cells in an intact cell layer (black, solid). Both *p*<0.001, with *n*=3932 and n=4368 for Neighbor and Control, respectively. (b) AFM topography height scan shows continuous junctions and homogenous structures across the cell body in K14 rescue cells. (c,d) Immunostaining for α-Tubulin, ZO1, Claudin4 and E-cadherin in WT, Kty$^{-/-}$ and K14 cells 48h after $Ca^{2+}$-switch. Scale bars: 10 μm. (e) WB of total protein lysates for Occludin, Claudin1, Claudin4 and Tubulin. (e') Quantification of protein levels relative to tubulin as Loading control (mean+/-SEM, n=8).

### Videos

Video SI1: Wound closure in WT cells

Video SI2: Wound closure in $Kty^{-/-}$ cells

Video SI3: Wounding process in $Kty^{-/-}$ cells

Video SI4: Wounding process in WT cells

Video SI5: Wound closure in K14 cells

Video SI6: Wounding process in K14 cells
